# Imputation of canine genotype array data using 365 whole-genome sequences improves power of genome-wide association studies

**DOI:** 10.1101/540559

**Authors:** Jessica J. Hayward, Michelle E. White, Michael Boyle, Laura M. Shannon, Margret L. Casal, Marta G. Castelhano, Sharon A. Center, Vicki N. Meyers-Wallen, Kenneth W. Simpson, Nathan B. Sutter, Rory J. Todhunter, Adam R. Boyko

## Abstract

Genomic resources for the domestic dog have improved with the widespread adoption of a 173k SNP array platform and updated reference genome. SNP arrays of this density are sufficient for detecting genetic associations within breeds but are underpowered for finding associations across multiple breeds or in mixed-breed dogs, where linkage disequilibrium rapidly decays between markers, even though such studies would hold particular promise for mapping complex diseases and traits. Here we introduce an imputation reference panel, consisting of 365 diverse, whole-genome sequenced dogs and wolves, which increases the number of markers that can be queried in genome-wide association studies approximately 130-fold. Using previously genotyped dogs, we show the utility of this reference panel in identifying novel associations and fine-mapping for canine body size and blood phenotypes, even when causal loci are not in strong linkage disequilibrium with any single array marker. This reference panel resource will improve future genome-wide association studies for canine complex diseases and other phenotypes.

**Author Summary:** Complex traits are controlled by more than one gene and as such are difficult to map. For complex trait mapping in the domestic dog, researchers use the current array of 173,000 variants, with only minimal success. Here, we use a method called imputation to increase the number of variants – from 173,000 to 24 million – that can be queried in canine association studies. We use sequence data from the whole genomes of 365 dogs and wolves to accurately predict variants, in a separate cohort of dogs, that are not present on the array. Using dog body size, we show that the increase in variants results in an increase in mapping power, through the identification of new associations and the narrowing of regions of interest. This imputation panel is particularly important because of its usefulness in improving complex trait mapping in the dog, which has significant implications for discovery of variants in humans with similar diseases.

## Introduction

The modern domestic dog (*Canis lupus familiaris*) consists of over 500 breeds selected for diverse roles and subject to wildly different disease prevalences (1). A high quality reference genome (2–4) and affordable SNP genotyping arrays (5) have helped make the dog a powerful animal model for studying the genetics of complex traits and diseases. Of 719 traits genetic traits and disorders in the dog, 420 are potential models of human disease (https://omi.org/home/). With an average spacing of 1 SNP every 13kb, the CanineHD array (Illumina, San Diego, CA) has been successfully implemented in many genome-wide association studies (GWAS), especially within single breeds where linkage disequilibrium (LD) often extends beyond 1Mb (for example, see (6,7)). However, the results of many complex disease mapping studies in dogs have been underwhelming, with only one or a few significant loci identified (for example, see (8–10)). 57% of the 719 genetic traits and disorders in dogs are complex but the likely causal variant is known for only 27% of these (https://omia.org/home/).

Recently, we used simulations to show that an increase in SNP density to 1 SNP every 2kb would improve power for canine complex trait GWAS (8). An increase in density can be achieved by the following: adding more SNPs to the CanineHD array, using whole genome sequencing (WGS), or using imputation to predict genotypes through the use of a reference panel created from WGS data. Of these, imputation is the most cost-effective option and has been used successfully in human and cattle GWAS, especially with the recent WGS efforts in these species (11,12). GWAS of canine morphological traits has been very successful, due to large effect sizes and long regions of LD as a result of recent selection in purebred dogs (13). Seventeen quantitative trait loci (QTLs) associated with body weight, as a proxy for body size, have been identified (5,8,14– 22), as well as associations for other morphological phenotypes such as ear flop (5,15,16) and fur type (8,23,24). Despite the success of morphological trait mapping, we suggest that imputation can improve the power of GWAS, especially for reducing large intervals for use in fine-mapping. We posit that improving the density of variants by using an imputation panel will greatly improve the power to identify causal loci for canine complex traits, due to increased LD. We use 365 canine whole genome sequences to create a reference panel of 24 million variants and impute these variants in 6,112 dogs previously genotyped on a semi-custom 185k CanineHD array. We show that using an imputation panel increases our power to detect variants affecting complex canine traits – both morphological and blood phenotypes – by identifying novel loci, and by refining intervals for previously-identified QTL’s for use in fine-mapping. To our knowledge, this is the first study to use an imputation panel based on WGS for canine mapping studies.

## Results

### Evaluation of imputation accuracy

We used IMPUTE2 to impute the WGS reference panel across the 6,112 genotyped dogs resulting in 24 million variants. By comparing 33,144 imputed variants to directly genotyped sites on a second custom array, we were able to calculate the accuracy for our imputation panel, which was 88.4% overall. Across all sites, purebred dogs had the highest accuracy (89.7%, n=276), followed by mixed-breed dogs (88.6%, n=13), and then village dogs (84.2%, n=86). This result is expected given that 210 of our 365 WGS panel were purebred dogs, and also due to the long-range haplotypes found in purebreds that make calling imputed variants easier. For all three dog types (purebred, mixed-breed, and village), imputation accuracy increased with decreasing minor allele frequency (MAF) (Fig. 1a), which is an expected result because as MAF decreases, the occurrence of the major allele is the correct call more often. Looking at true heterozygous sites only (Fig. 1b), imputation accuracy was lower across all MAFs compared to all sites (Fig. 1a). Imputation accuracy increased as MAF increased for heterozygous sites, as there are more heterozygous calls for SNPs with higher MAF.

**Figure 1:**
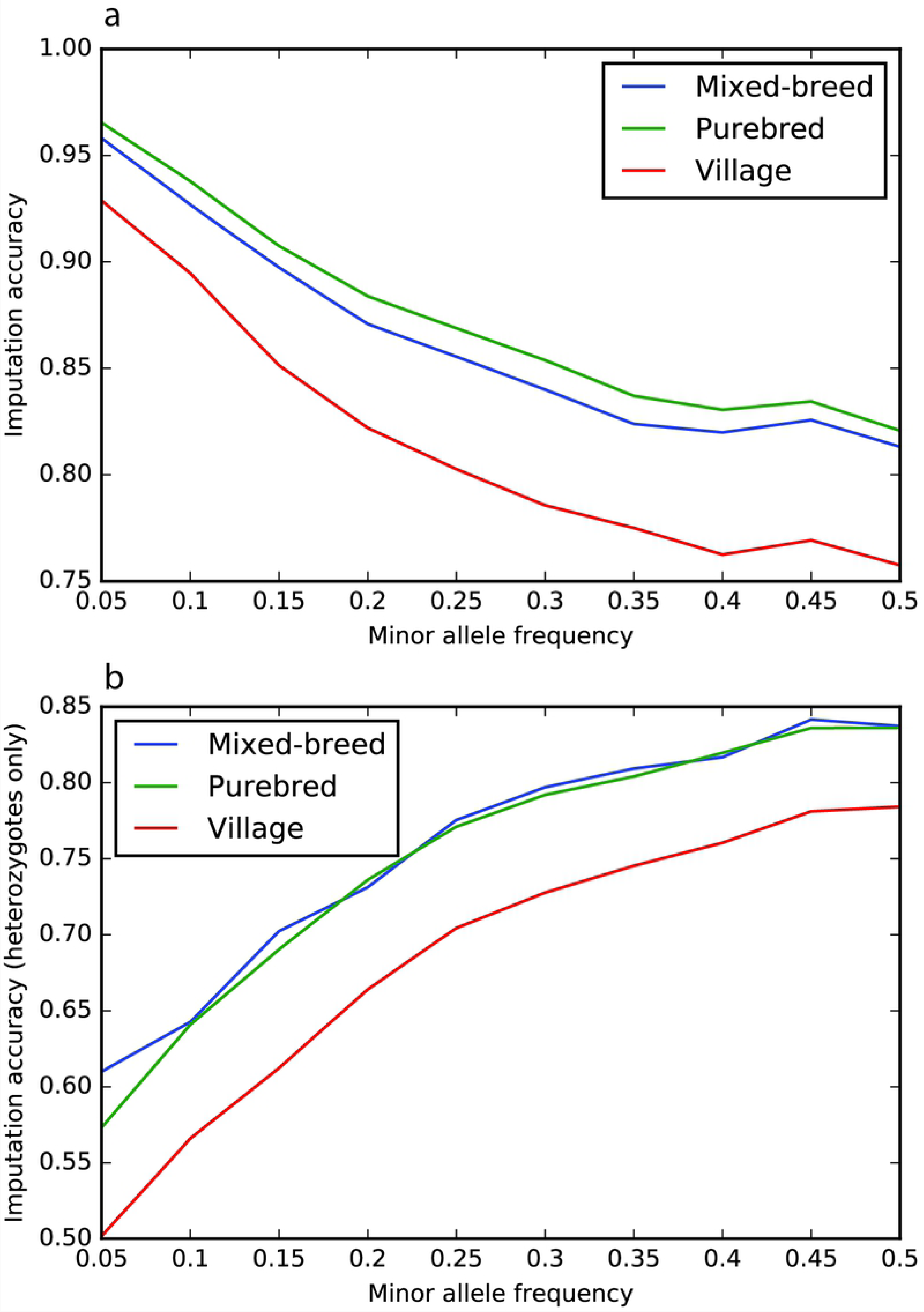
Imputation accuracy for 33,144 variants of different allele frequency in mixed-breed dogs (blue), purebred dogs (green), and village dogs (red). a) all sites, b) heterozygous sites only.

In general, the larger chromosomes and chromosome X had higher imputation accuracies than the smaller chromosomes, such as 35, 36, 37, and 38, due to lower recombination rates (S1 Table). For all purebred, mixed-breed, and village dogs, the average imputation accuracy per chromosome was 89.1% (range of 84.3-93.5%), 88.0% (range of 83.0-93.0%), and 83.7% (range of 78.5-92.4%), respectively.

### Body size associations

We performed two separate GWAS, firstly using the semi-custom CanineHD array data of 185k markers, and secondly using our imputed panel of 24 million variants. The phenotypes used in these GWAS were male breed-average weight^0.303^, male breed-average height, and individual sex-corrected weight^0.303^. We were then able to compare the results from the array GWAS and the imputed GWAS using the exact same phenotypes.

Using imputed data generally increased the significance of body size associations seen in the array data, especially *HMGA2* on CFA10 and fgf4 on CFA18 for height (Fig 2a,b,c; S2 Table). Most of the respective QTLs from the array GWAS and the imputed GWAS were in LD (r^2^ > 0.2) with the exception of the CFA12 and two CFA26 associations (Table 1). When imputed variants were not in high LD (r^2^ < 0.8) with array QTLs, the imputed variants generally had stronger effect sizes and lower minor allele frequencies (Table 1).

**Table 1:**
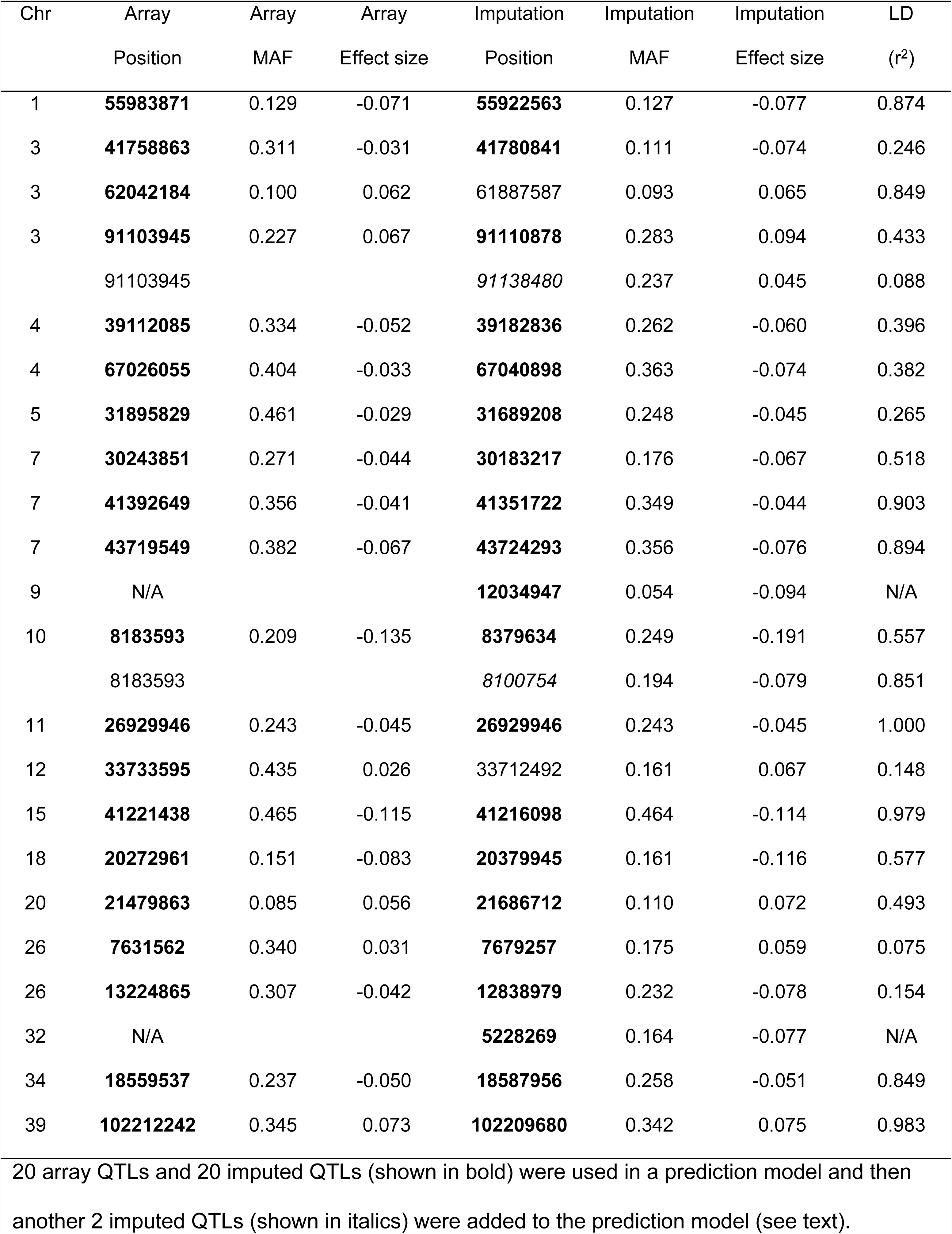
Positions, minor allele frequencies (MAF), and effect sizes for SNPs associated with breed-average male weight0.303 using the array data and imputed data, and LD between these pairs of SNPs.

**Figure 2:**
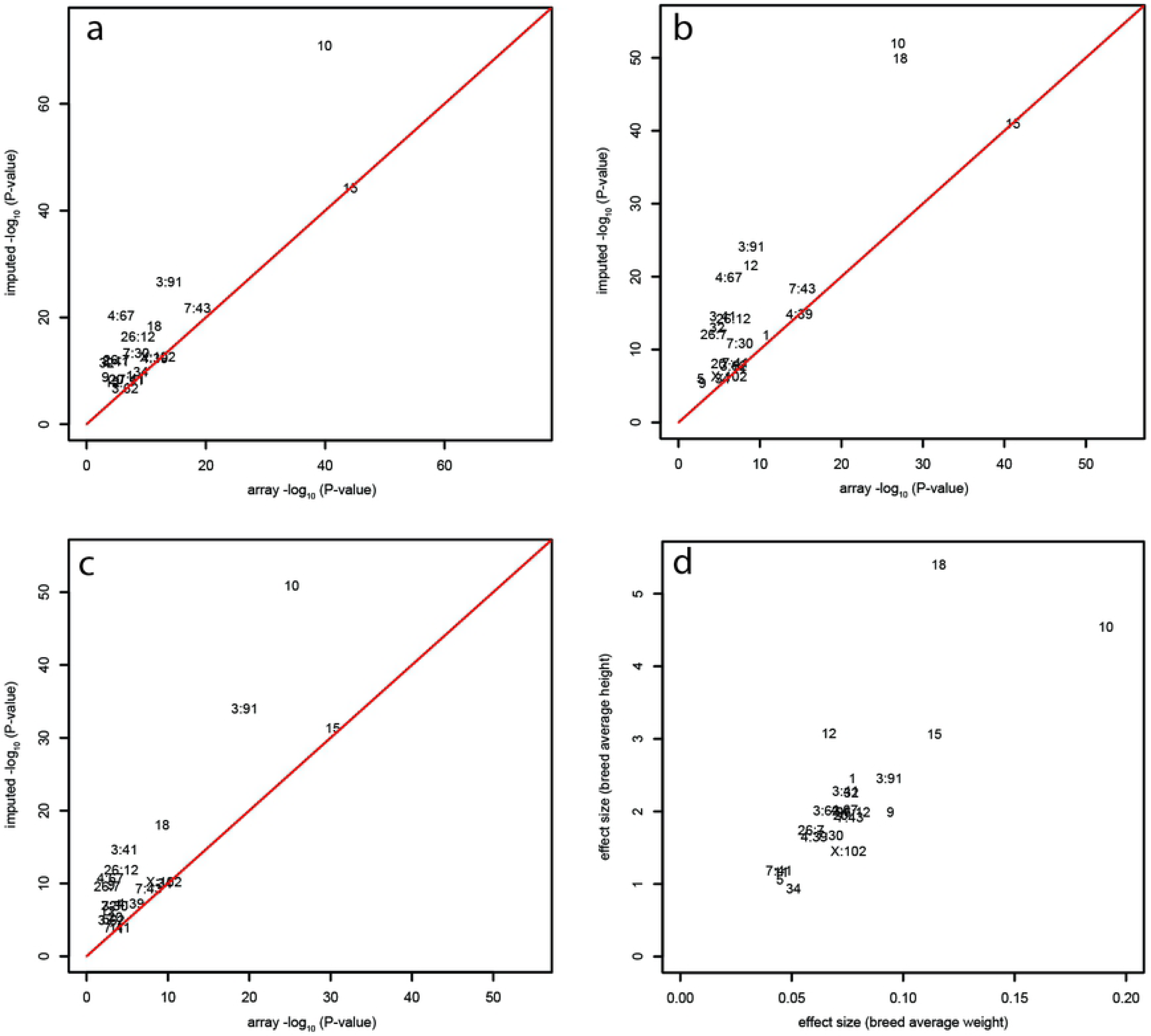
Scatter plots showing *P*-values for array GWAS (x axis) and imputed GWAS (y axis). a) breed-average weight, b) breed-average height, c) individual weight, and d) absolute values of the effect sizes of the most-associated SNP for breed-average weight and breed-average height from the imputed GWAS.

For most size-associated variants, the breed-average weight and height effects were roughly isometric, with the exception of CFA18 and CFA12 QTLs, which had a greater effect on height than weight (Fig. 2d). Of the seventeen autosomal QTLs that have been previously associated with body size (5,8,14–22,25), only one (CFA3:62) did not reach significance using the imputed panel (significance threshold of *P* = 1×10^−8^), with *P*-values of 1.1×10^−5^, 2.1×10^−7^ and 1.8×10^−8^ for individual weight, breed-average weight and breed-average height respectively (S2 Table). GWAS of breed-average weight provided the most power on average, so we will focus on that phenotype for the rest of the body size analyses.

We used the identified body size QTLs to predict the body weight of individuals, by randomly setting 20% of the body weights in the dataset to missing, then using a Bayesian sparse linear mixed model to predict the missing weights, and finally comparing the predicted weights to the actual weights. Using the 20 QTLs identified from the array GWAS (see bold in Table 1), we found a correlation coefficient (r) of 0.851. Using the 20 QTLs from the imputed GWAS (see bold in Table 1), we found r of 0.869, and this increased very slightly to 0.870 when two more QTLs were included (see italics in Table 1).

### Known body size loci

Nine body size QTLs have previously been fine-mapped (*IGF1R, STC2, GHR, SMAD2, HMGA2, FGF4, IGF1*, fgf4, *IGSF1*) (17–20,25,26). For each of these, the region in LD with the most significant respective marker in the breed-average weight imputed GWAS contained the known or putative causal variant (S1 Fig.). While the putative causal variants weren’t always the highest associated variant at a locus, they generally had *P*-values within two orders of magnitude of the most associated marker (S1 Fig.), confirming that the imputation panel performs well for the weight GWAS.

Unsurprisingly, many of the putative causal variants are not markers on the CanineHD array, including *IGF1R* (3:41,849,479) (20), *STC2* (4:39,182,836), *GHR* (4:67,040,898 and 4:67,040,939), and *HMGA2* (10:8,348,804) (17). With the array data, the *IGF1R* QTL (*P* = 1.4×10^−5^ for breed-average weight GWAS) did not reach significance, but with the imputed data we saw a significant association signal (*P* = 1.6×10^−12^ for breed-average weight GWAS), and the causal variant was the 6^th^ most associated SNP (r^2^ = 0.73 between causal and associated SNP) (S1 Fig. a). The *STC2* and *GHR* (4:67,040,898) putative causal variants were the most significant variants at those loci in the imputed GWAS (S1 Fig. b,c). Note that there are two putative causal variants for *GHR* (17), both in exon 5, but only one passed our 5% MAF filter. Similarly, the *HMGA2* causal variant was in high LD (r^2^=0.91) with the most significant marker at this locus in the imputed GWAS (S1 Fig. e).

For *IGF1*, SNP5 (BICF2P971192, 15:41,221,438), which is in LD with the SINE element (18), was the most significant association in the array GWAS. In the imputed GWAS, SNP5 was the 2^nd^ most associated SNP and the SNP that tags the SINE element (15:41,220,982) was the 4^th^ most associated SNP, and these SNPs were nearly in complete LD with the most significant marker in the GWAS (r^2^=0.98 and 0.97 respectively) (S1 Fig. f). The *IGSF1* missense mutation (26) was in high LD with the most significant association in the imputed GWAS (r^2^=0.97) (S1 Fig. i). Note that there is a second variant in *IGSF1* – an in-frame deletion – that has also been identified (26).

### Imputed GWAS novel body size loci

We previously identified four novel QTLs, making a total of seventeen, that are associated with body size in dogs (8). Here, using the imputation data, we found a further five novel QTLs (CFA5:31, CFA7:41, CFA9:12, CFA26:7, CFA32:5) that passed our significance threshold in a breed-average weight GWAS. As a conservative control, we performed a further breed-average weight GWAS in which we included the four most-associated QTLs (CFA10:8, CFA15:41, CFA3:91, CFA7:43) as covariates. The results showed that the novel loci at CFA5:31, CFA7:41 and CFA32:5 were no longer significant – and four other loci were also not significant in the covariate GWAS: CFA1:56, CFA11:26, CFA20:21, CFA34:18. Further analyses are required to determine if these are true or spurious associations, but since we cannot rule out that they are spurious, we conclude that we have identified only two novel canine body size QTLs, at CFA9:12 and CFA26:7.

The most significant SNP at CFA9:12 is located about 200kb upstream of the gene *growth hormone 1* (*GH1)* (Fig. 3a), which is expressed in the pituitary and has been associated with body size in humans and cattle (27–29). The non-reference, derived indel that was the most highly associated in our imputed GWAS is found at high frequency in two small breeds Papillon and Pomeranian, and also in New Guinea Singing Dogs. Using our snpEff annotated variant files, we found two variants in *GH1*: a splice donor variant in intron 3 (CFA9:11,833,343, c.288+2_288+3insT), and an in-frame deletion in exon 5 (CFA9:11,832,437, c.573_578delGAAAGA, p.Lys191_Asp). Both of these variants were at <5% frequency in the WGS panel but all occurrences were in small-sized breeds (such as Yorkshire terrier and Maltese).

**Figure 3:**
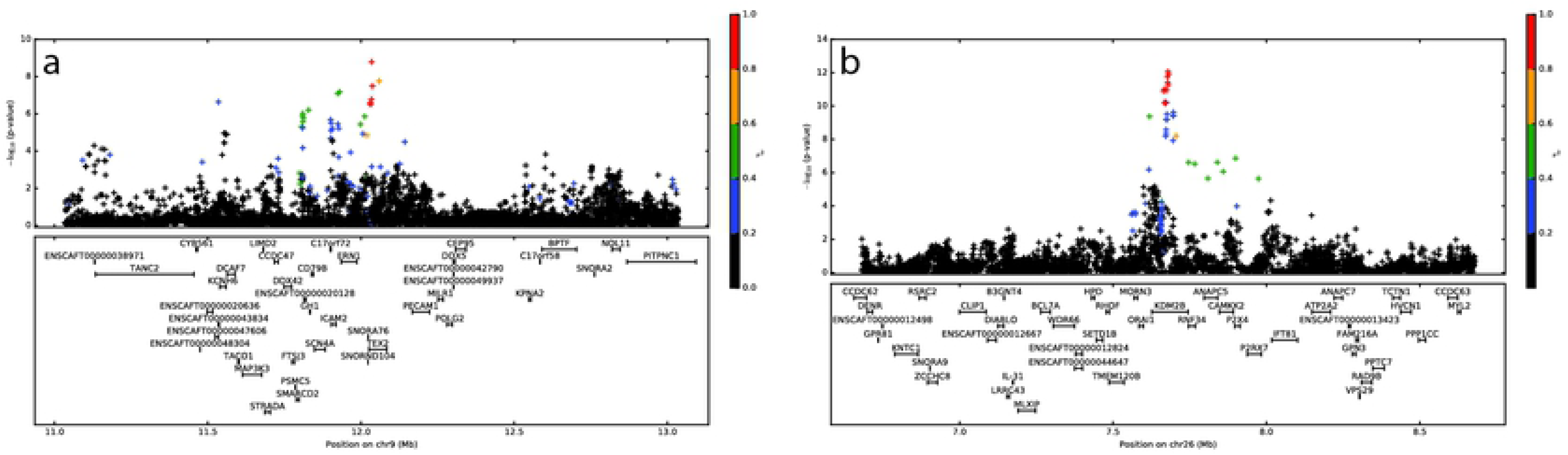
Novel body size loci.Linkage disequilibrium plots of the region around breed-average weight GWAS results a) CFA9:12, b) CFA26:7. Array genotypes are shown as o, imputed data are shown as +. Colors indicate amount of LD with the most significantly associated SNP, ranging from black (r^2^<0.2) to red (r^2^>0.8).

The second novel body size QTL is at CFA26:7 (Fig. 3b). Investigation of the surrounding region uncovered a couple of potential candidate genes. The first is *ANAPC5*, a member of the anaphase-promoting complex gene family that includes *ANAPC13*, which has been associated with height in humans (30). The second candidate gene is the histone H3 demethylase *KDM2B* (*Iysine-specific demethylase 2B)*, which has been associated with body mass index in humans in a CpG methylation study (31). However, we did not identify any variants in *ANAPC5* or *KDM2B* in the snpEff-annotated files that are in LD with the associated imputation variant. The non-reference, derived allele was found at high frequency in the small breeds Shiba Inu, Havanese, and Chihuahua.

### Refinement of body size loci

With the imputation panel, we saw a refinement in several QTL regions – for example, the chromosome 3 association near the genes *LCORL* and *ANAPC13,* both of which have previously been associated with body size (5,15,29,30,32). Using imputed data, this QTL had a more significant and defined association, compared to the CanineHD array data alone (S2 Fig. a). The QTL interval is about 65kb and 60kb upstream of the genes *LCORL* and *ANAPC13* respectively, suggesting the causal variant is likely regulatory. Another example is the recently identified body size QTL at CFA7:30 Mb, near the gene *TBX19* (8). Here the imputed GWAS results showed a narrower QTL interval of greater significance when compared to the array GWAS (S2 Fig. b). This region overlaps *TBX19* but we did not observe any coding loci in our snpEff annotated variant files that are in LD with the most associated SNP.

### Allelic heterogeneity

In order to reduce phenotypic noise, again we included the four most-associated QTLs (CFA10:8, CFA15:41, CFA3:91, CFA7:43) as covariates in the GWAS (hereafter referred to as “top 4 covariates”), and then implemented a region-specific stepwise approach, including further associated SNPs in the region as covariates, until no significant association signal remained. For breed-average body weight, when we regressed out the most significant association for a QTL, we expected the association signal to disappear, as seen with the *SMAD2* QTL (Fig. 4a). Our results showed two QTLs (CFA3:91, CFA10:8) that retain significant association signal after regressing out the most associated locus in the respective region (Fig. 4b,c). For both CFA3 and CFA10, the data suggest there may be two independent significant associations in these regions. In the CFA3 region, the initial association signal peak looked regulatory while the residual signal is located in the genes *ANAPC13* and *LCORL*. The residual signal in the CFA10 region lies close to the ear flop association (5,15,16) (candidate gene *MSRB3*) but is not in LD with the imputed ear flop locus at CFA10: 8,097,650 (r^2^=0.147). The variant in this residual signal region may be regulatory, as the significance peak lies approximately 248kb upstream of *HMGA2*. We did not identify any coding variants in these two residual signal regions from the snpEff annotations. Two other QTLs (CFA4:67 and CFA15:41) showed evidence of residual signal but these did not reach significance (*P* = 2.9×10^−7^ and 2.9×10^−8^, respectively).

**Figure 4:**
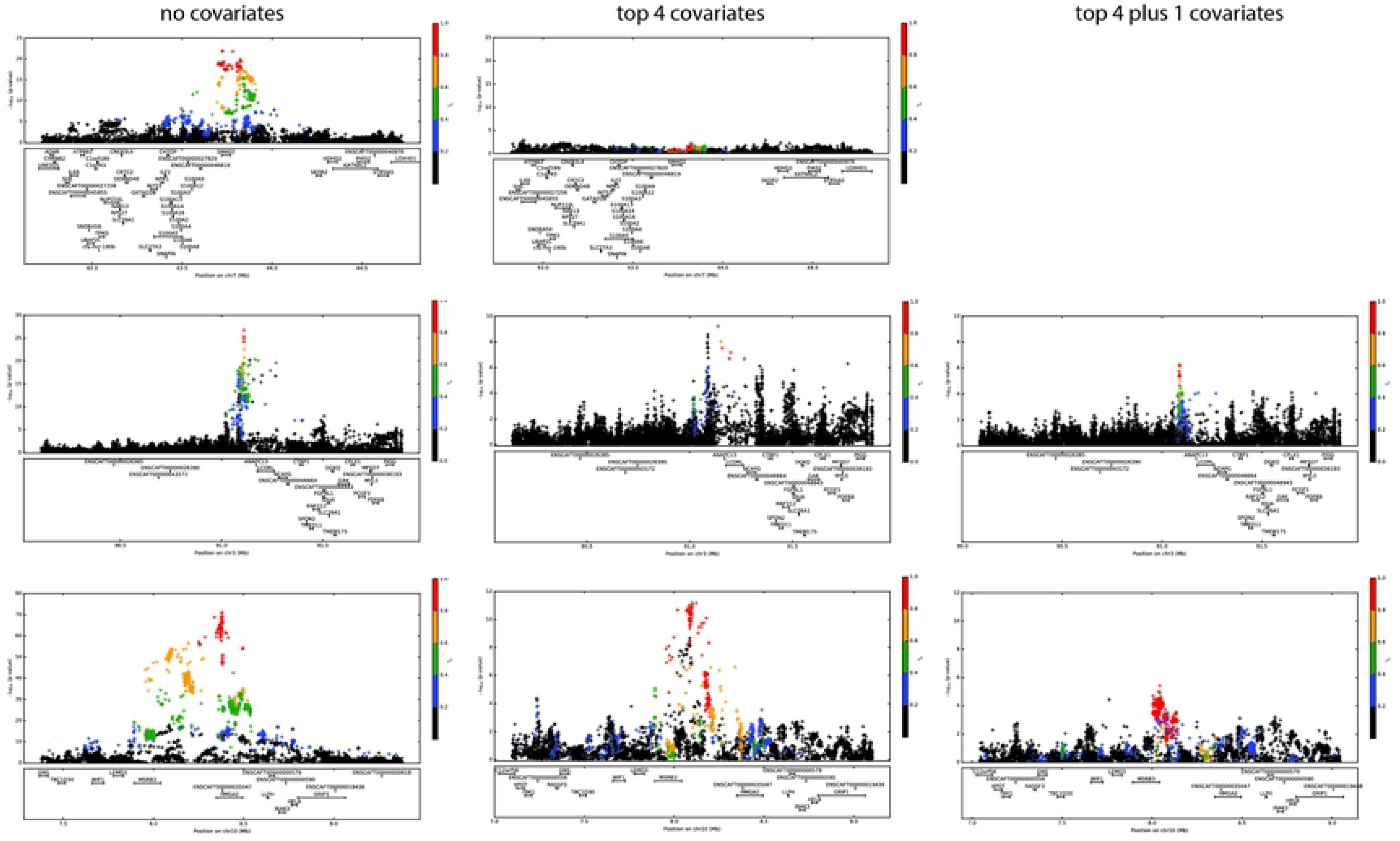
Stepwise-covariate LD plots for breed-average weight imputed GWAS. a) CFA7 (*SMAD2*), showing that the association signal in the region disappears with the inclusion of the top 4 SNPs as covariates. b)-d) Association signal in the region remains with the inclusion of the top 4 SNPs as covariates and then additional stepwise covariates in the region b) *LCORL*, c) *HMGA2*. Array genotypes are shown as o, imputed data are shown as +. Colors indicate amount of LD with the most significantly associated SNP, ranging from black (r^2^<0.2) to red (r^2^>0.8).

This residual signal suggested allelic heterogeneity in these regions but could also be due to imperfect tagging in the imputed dataset. As a follow-up analysis, for each of these two QTLs (CFA3:91 and CFA10:8), we took the most significant SNP from the top 4 covariates GWAS. We used that significant SNP as a covariate in a GWAS to see if we were able to recover the most significant SNP from the initial GWAS with no covariates. For both CFA3 and CFA10, we did recover the initial associated SNP, suggesting that these are real associations and not midway between two imperfectly tagged SNPs.

### Blood phenotypes

Using our imputed panel for GWAS on blood phenotypes revealed several novel associations. For example, we saw significant associations with the phenotypes of albumin and calcium levels in peripheral blood (*P* = 4.5×10^−10^ and 5.9×10^−9^ respectively), neither of which were previously identified in the array GWAS (33) (S3 Table). We also identified a novel association with blood glucose level and CFA1, located in the gene solute carrier family 22 member 1 (S*LC22A1)* and about 30kb downstream of the insulin-like growth factor 2 receptor gene *(CI-MPR/IGF2R)* (Fig. 5d). During gestation, *IGF2R* binds insulin-like growth factor 2 (IGF2), the presence of which stimulates the uptake of glucose (34). The SNP was at highest frequency (>50%) in the Samoyed and American Eskimo dog breeds.

**Figure 5:**
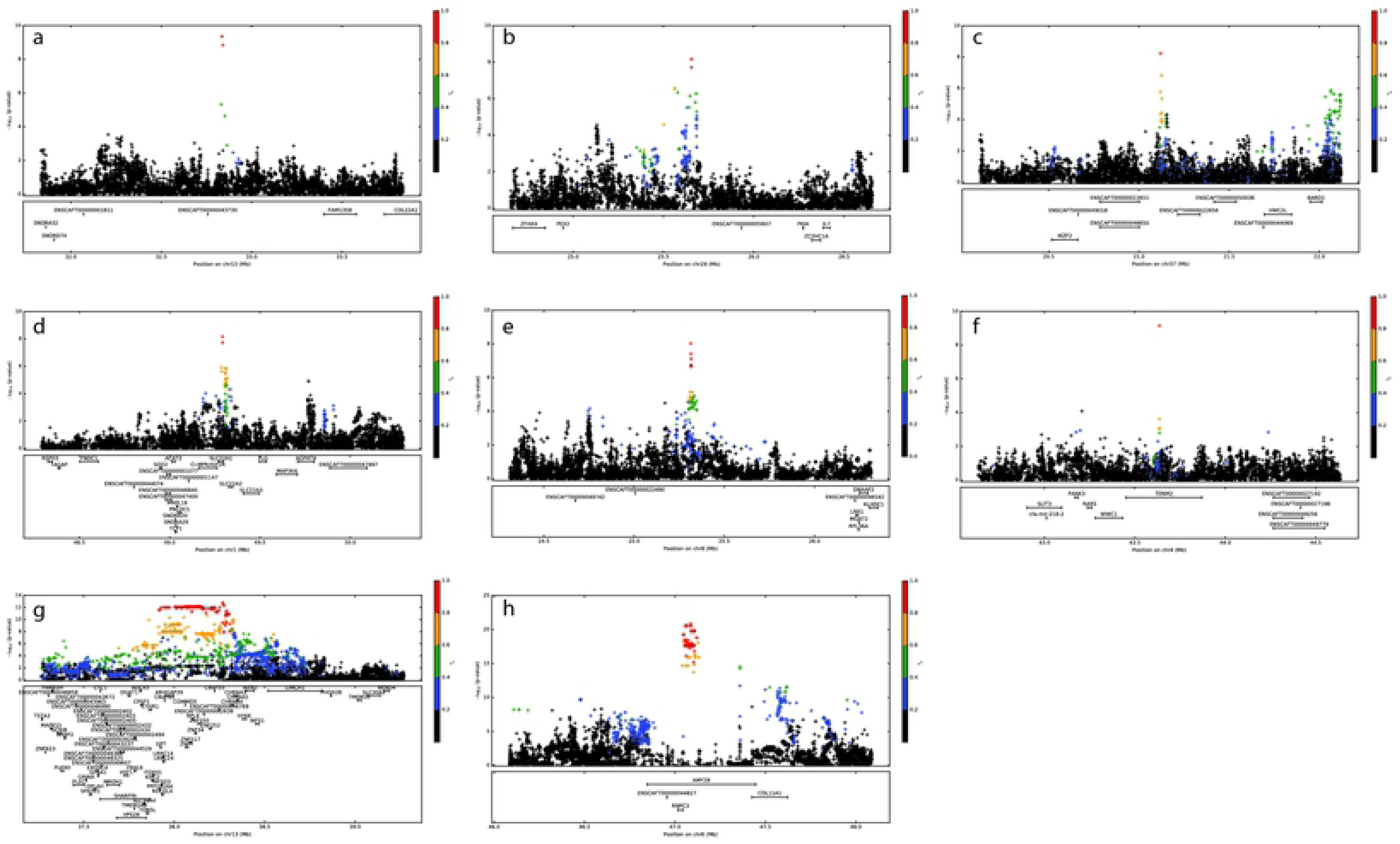
LD plots for significant blood phenotypes. a) albumin on CFA13, b) anion gap on CFA29, c) calcium on CFA37, d) sqrt glucose on CFA1, e) potassium on CFA8, f) white blood cells on CFA4, g) log ALT on CFA13, h) sqrt amylase on CFA6. Array genotypes are shown as o, imputed data are shown as +. Colors indicate amount of LD with the most significantly associated SNP, ranging from black (r^2^<0.2) to red (r^2^>0.8).

Of the eight significant associations (using a threshold of *P* = 1.0×10^−8^) we saw with the imputed data, only two were not novel – alanine aminotransferase (ALT) and amylase – although both increased in significance (Fig. 5g, 5h; S3 Table). In addition to significant associations, we also saw six associations that nearly meet our significance threshold, that is, *P* < 2.0×10^−8^, including three that were not significant using the genotype data (S3 Table).

## Discussion

Imputation increases GWAS power by including additional sites that are not well-tagged by any single array marker, and has been successfully implemented in human studies, for example, low-density lipoprotein GWAS (35–37). Here we present a canine imputation panel of 24 million variants – an approximate 130-fold increase in SNP number and SNP density from the semi-custom CanineHD array – for use in association studies. This panel has an overall accuracy rate of 88.4% when compared to genotype data from the same individuals (276 purebreds, 86 village dogs, and 13 mixed-breed dogs). In the future, panels based on even larger numbers of sequenced individuals would yield even higher accuracy (for example, see (38)), but this panel based on hundreds of dogs is still a useful, cost-effective way to improve the power of canine mapping studies today.

With our imputation panel, we improved association mapping for previously studied phenotypes, such as body size. Previous mapping studies of canine body size and other morphological traits using CanineHD array data have identified many significant QTLs. This success is largely the result of selection for body size during the formation of dog breeds, leading to selective sweeps around large-effect loci that facilitated mapping efforts. Nevertheless, using the imputation panel, we were able to identify two additional novel loci (at CFA9:12 and CFA26:7) that influence body size although functional studies, which are beyond the scope of this research study, are required to validate these two loci. Using imputation, we were also able to narrow intervals for previously known associated QTLs, and find evidence of possible allelic heterogeneity at two loci. Furthermore, imputation provides a more accurate analysis of the genetic architecture underlying canine body size and, in turn, allows a more accurate prediction for body size in dogs. Imputation is especially helpful in across-breed and/or mixed-breed study designs, where LD breaks down very rapidly making it more difficult to identify associations. Increasing the number and density of queried variants (as done by imputation) increases the chance that a variant will be in LD with the causal variant, especially when compared to a within-breed study design. We used our imputation panel for across-breed GWAS of blood phenotypes, resulting in several novel associations and the narrowing of associated intervals when compared to array data alone. Although costs of WGS are decreasing, it is still more cost-effective to use a panel of WGS individuals to create an imputation dataset based on genotyped samples than it is to directly WGS all the samples (39). Our imputation panel was created using over 350 canine WGS’s representing 76 breeds. The inclusion of more breeds, especially diverse breeds (such as the Parson Russell Terrier) and rare breeds (such as the Pumi), will improve the accuracy of, and the number of rare variants in, future imputation panels. A recent canine imputation study has shown that imputation accuracy is highest using a multi-breed reference panel (compared to a breed-specific panel), and when there is overlap in breeds between the target and reference panel (40). Furthermore, human studies have shown that imputation accuracy increases with the size of the reference panel (41,42).

In summary, using our canine imputation panel of 24 million variants results in an increase in GWAS power, even for phenotypes that have multiple significant associations. The improvements to canine GWAS, especially for complex phenotypes, will not only further the field of canine genetics, but may also have beneficial implications for human medical genetics – especially for complex diseases, such as cancer, for which the domestic dog is a good model organism (43).

## Material and Methods

### Whole genome sequences

The 365 whole genome sequences include 210 breed dogs (from 76 breeds), 107 village dogs (from 13 countries), and 28 wolves (S4 Table). 88 of these were sequenced at the Cornell University BRC Genomics Facility; others were sourced from public databases (S4 Table). Those sequenced at Cornell were run on an Illumina HiSeq2000 or Illumina HiSeq2500 and the reads were aligned to the CanFam3.1 reference genome using BWA (44). Variants were called using GATK’s HaplotypeCaller (46–48). Variant quality recalibration was done in GATK using the semi-custom canineHD variant sites (8) as a training set (known=false, training=true, truth=true, prior=12.0). We included SNPs in the 99.9% tranche and removed sites with minor allele frequency (MAF) less than 0.5%. Phasing was done using Beagle r1399 (45).

### Imputation panel

SHAPEIT v2.r790(49) was used to phase the genotype data from 6,112 dogs as previously described (8) and then IMPUTE2 version 2.3.0 (50) was used to impute across these data. Imputation was only performed on the autosomes and chromosome X, not on the Y chromosome or mitochondrial SNPs. The final reference panel consists of 24.0 million variants, including 750,000 on the X chromosome, of which 20.33 million are SNPs and 3.67 million are indels.

### Imputation Accuracy

276 purebred, 86 village, and 13 mixed-breed dogs were also genotyped on a second custom Illumina CanineHD 215k array, which contains 33,144 SNPs that are not on the 185k semi-custom CanineHD array but do feature in our imputed dataset. These 33,144 SNPs were used to determine the accuracy of our imputation across the 375 total dogs. Accuracy was calculated for each SNP as the number of sites that are correctly called in the imputed dataset divided by the total number of dogs. For example, if a G/C SNP was called G/G in 14 dogs, C/C in 10 dogs, and G/C in the remaining 351 dogs, then the imputation accuracy for that SNP is 93.6%. MAF was calculated for each SNP as the number of occurrences of the allele across all village, purebred, and mixed-breed dogs in the genotyped dataset. Imputation accuracy and MAF were plotted in Jupyter notebook (51) using Matplotlib library (52).

### Marker Datasets

For the genotype data, individuals were run on a semi-custom Illumina CanineHD array of 185k SNPs, and quality control steps were performed as previously described(8). For the imputed data, IMPUTE2 outputs were converted into PLINK (53) binary format, one for each chromosome.

### GWAS

We ran GWAS using a linear mixed model in the program GEMMA v 0.94 (54), with a MAF cut-off of 5% and using the Wald test to determine *P*-values. All LD plots were created using Matplotlib library (52) in Jupyter notebook (51).

For the imputed panel, GWAS was performed for each canine chromosome (CFA1-39) separately. The kinship matrix calculated using the array data in GEMMA was also used in the imputed GWAS for the same phenotype. The significance threshold was set to *P* = 1×10^−8^ (see (55)).

#### 1. Body size

To identify QTLs associated with body size, we ran a GWAS of breed-average body weights (n=1926) and heights (n=1926), and individual body weights (n=3095), using both the semi-custom 185k CanineHD array data and the imputed panel data. Effect sizes were recorded from the GEMMA output, and MAF’s and LD statistics were calculated using PLINK.

##### Breed-average body size

The breed-average data included dogs from 175 breeds with a maximum of 25 dogs per breed. The phenotypes of male breed-average weight^0.303^ in kg, or male breed-average height in cm, were assigned to all dogs in the breed for the weight and height GWAS, respectively. We used weight^0.303^ to normalize the distribution of weights across the breeds, based on a Box-Cox transformation performed in R (56) using the package MASS (57).

##### Individual body weights

Individual body weight GWAS was performed using 3,095 dogs including 417 village dogs and 427 mixed-breed dogs. The average raw weight in this individual dataset is 24.7kg, ranging from 1.6kg to 99.7kg. 164 breeds are represented, with 52 small breeds, 57 medium breeds, 34 large, and 21 giant breeds. Individual weights were sex-corrected by 16.47% (that is, female weights were increased by 16.47%), and transformed (weight^0.303^).

##### Covariate GWAS

GWAS of breed-average body weight using imputed variants shows the four most-associated loci are *HMGA2* (CFA10), *IGF1* (CFA15), *LCORL* (CFA3), and *SMAD2* (CFA7). In order to control for possible spurious associations, we ran an imputed GWAS, using breed-average body weights, including the four most-associated loci as covariates and then observed which of our significant associations remained.

##### Fine mapping

We used snpEff version 4.3T (58) to annotate our WGS variant file, using the pre-built CanFam3.1.86 database, and then used this to search for potential causal variants within specific LD regions.

##### Predicting body size

We used all the individual body weights (n=3,095) and specific body size QTLs as a training set, and then randomly set 20% of the weights to missing and used a Bayesian sparse linear mixed model (with a ridge regression/GBLUP fit) in GEMMA to predict these missing weights. We did this randomization and prediction 50 times, and then compared the actual weights to the predicted weights using a correlation coefficient.

##### Allelic heterogeneity

In order to reduce phenotypic noise, we again included all four most-associated QTLs as covariates in a follow-up GWAS of breed-average body weight and then, for those regions that still had residual association signal, we also included the most associated SNP (in addition to the four) as a covariate in the next GWAS. We continued this stepwise process of including the most-associated SNP in the next GWAS until there was no significant association signal in the respective region remaining. The same analysis was run using the three most-associated loci (*HMGA2, IGF1, LCORL*) and the five most-associated loci (*HMGA2, IGF1, LCORL, SMAD2, GHR*) with comparable results (data not shown).

#### 2. Blood phenotypes

Using previously published data of 38 phenotypes from complete blood count (CBC) and serum chemistry diagnostic panels (33), GWAS were performed using the imputed data and results were compared to the published results using the semi-custom CanineHD array data.

### Data Availability

Whole-genome sequence data produced for this research have been deposited in the Sequence Read Archive (http://www.ncbi.nlm.nih.gov/sra) with accession numbers listed in S4 Table. The pre-phased genotype data (binary PLINK files) and the phased WGS data (vcf files) are publicly available at datadryad.org (doi:10.5061/dryad.jk9504s).

## Acknowledgments

The authors would like to acknowledge Liz Corey and other Cornell Veterinary Biobank personnel, Peter Schweitzer at the Cornell University Genomics Facility, and the many dog owners that contributed samples.

## Author Contributions

Investigation: JJH, ARB

Formal analysis: MB, LMS

Resources: MEW, MLC, MGC, SAC, VM-W, KWS, NBS, RJT

Writing – original draft preparation: JJH, ARB

Writing – review and editing: JJH, MEW, MB, LMS, MLC, MGC, SAC, VM-W, KWS, NBS, RJT, ARB

## Supporting Information

S1 Figure: Linkage Disequilibrium (LD) plots of the region around the breed-average weight QTL intervals that have been fine-mapped a) CFA3:41 near the gene *IGF1R*, b) CFA4:39 near the gene *STC2*, c) CFA4:67 near the gene *GHR*, d) CFA7:43 near the gene *SMAD2*, e) CFA10:8 near the gene *HMGA2*, f) CFA15:41 near the gene *IGF1*, g) CFA18:20, h) CFA32:5 near the gene *BMP3*, i) CFAX near the gene *IGSF1*. Dashed lines show the significant interval (defined by *P*-values within two orders of magnitude of the most associated SNP). Asterisks show the location of the causal locus. Note that for d), f), and g) the causal locus is a deletion, SINE insertion, and retrogene insertion respectively, so these locations are labeled with asterisks at the top of the plot.

S2 Figure: Linkage Disequilibrium (LD) plots of the region around the breed-average weight QTL intervals on a) CFA3:91 near the gene *LCORL* and *ANAPC13*, b) CFA7:30 near the gene *TBX19*. The significant interval, drawn with dashed vertical lines, is defined by *P*-values within two orders of magnitude of the most associated SNP. Array genotypes are shown as o, imputed data are shown as +. Arrows point to the most significant SNP in the array GWAS and imputed GWAS. Colors indicate amount of LD with the most significantly associated SNP, ranging from black (r^2^<0.2) to red (r^2^>0.8).

S1 Table: Imputation accuracy calculated for each chromosome for village dogs, mixed-breed dogs, and purebred dogs. Average recombination rate (cM/Mb) calculated from (59) for each chromosome is also shown.

S2 Table: *P*-values for body size GWAS QTLs using individual weight, breed-average weight, and breed-average height phenotypes with array genotypes and imputed panel. Novel body size loci are shown in bold.

S3 Table: Significant and nearly significant imputed GWAS results of blood phenotypes. Also shown is the location and *P*-value from the GWAS using the array genotype data.

S4 Table: List of Whole Genome Sequence (WGS) dog samples.

